# Genome-wide Identification of Transcriptional Start Sites and Candidate Enhancers Regulating Worker Metamorphosis in *Apis mellifera*

**DOI:** 10.64898/2026.03.12.711487

**Authors:** Kouhei Toga, Kakeru Yokoi, Hidemasa Bono

## Abstract

Eusociality in bees represents a major evolutionary transition and understanding its molecular basis is fundamental for sociogenomic studies. Comparative genomics has revealed correlations between transcription factor binding site (TFBS) abundance and social complexity; however, when and where these TFBSs function in a eusocial context remains largely unclear. In this study, we performed cap analysis of gene expression (CAGE) during worker metamorphosis in the honeybee *Apis mellifera* to identify TFBSs within active enhancers and decipher the regulatory relationships between these enhancers and their target genes. We identified 17,349 transcription start sites (TSSs) and 842 candidate enhancers. Using CAGE, we identified five clusters based on expression patterns. Notably, genes associated with the canonical metamorphic regulators, *Broad complex* (*Br-c*) and *E93*, were found within specific clusters. By integrating the correlations between enhancer and TSS activities with motif enrichment analysis, we identified 15 transcription factor–enhancer–TSS regulatory relationships. Among these, *tramtrack* (*ttk*)-binding sites were identified in five enhancers associated with four target genes, including *Br-c*. The number of target genes regulated by *ttk* was the highest in our dataset. To examine whether this regulatory relationship is conserved across bee species with varying levels of sociality, we analyzed the sequence conservation of *ttk*-binding sites in *Br-c* enhancers and found that perfect sequence conservation of *ttk*-binding site was restricted to the *Apis* genus. The *ttk*-binding sites of other target genes exhibited the same *Apis*-specific conservation pattern. Our findings suggest that gene regulatory relationships during worker metamorphosis occur in a lineage-specific manner in the *Apis* genus.

**Significance:** Honeybees produce distinct castes—queens and workers—from genetically identical larvae via differences in gene regulation. Although enhancers have been computationally predicted, their actual activity during bee development has rarely been measured directly, and the CAGE technology has never been applied for this purpose. We identified active enhancers during worker metamorphosis and discovered that the transcription factor *ttk* may regulate *Br-c*, a key developmental gene. This study provides the first direct evidence of active enhancers and their regulatory roles in honeybee worker metamorphosis.

## Introduction

Understanding of the molecular basis of eusociality, which represents a major evolutionary transition, in bees has been the central goal of sociogenomic studies (Robinson et al. 2005). Comparative genomic analyses across bee species with different levels of sociality have revealed that the number of transcription factor binding sites (TFBSs) correlates with the degree of sociality (Kapheim et al. 2015; Shell et al. 2021). The complexity of eusociality also correlates with the expansion of gene families in bees (Shell et al. 2021). Numerous transcriptome comparisons between castes and across developmental stages have led to the identification of candidate genes associated with social phenotypes in bees (Evans and Wheeler 1999; Cameron et al. 2013; Warner et al. 2019; Sasaki et al. 2021; Séguret et al. 2021; Soares et al. 2021; Toga and Bono 2023; Yokoi et al. 2025). Collectively, these findings highlight the central role of gene regulation in the evolution of eusociality.

Changes in the expression levels of genes can be readily identified using transcriptomic analysis. However, the regulatory transcription factors driving these changes remain largely unidentified because most TFBSs within enhancers are inferred from sequence-based conservation rather than direct observation of activity (Kapheim et al. 2015; Shell et al. 2021), with the exception of few (Wojciechowski et al. 2018; Jones et al. 2024). Consequently, when and where the predicted TFBSs function in social contexts remains largely unclear.

Enhancer RNAs (eRNAs) are a class of non-coding RNAs that are bidirectionally transcribed from enhancer regions by RNA polymerase II (Kim et al. 2010; Sartorelli and Lauberth 2020; Li et al. 2023). Because, similar to mRNAs, eRNAs possess a 5′-cap structure, they can be detected using the cap analysis of gene expression (CAGE), a method originally developed for identifying transcription start sites (TSSs) (Kodzius et al. 2006). In CAGE data, eRNAs appear as bidirectional peaks, the signals of which overlap with those of enhancer-associated histone modifications, such as H3K27ac and H3K4me1 (Andersson et al. 2014). These features are indicators of enhancer activity. Because CAGE can simultaneously detect eRNAs and TSSs, it is particularly effective in elucidating the relationship between enhancer activity and gene expression.

Although chromatin modifications were compared between queens and workers at the larval stage (96 h after hatching) using H3K4me3, H3K27ac, and H3K36me3 in a previous study (Wojciechowski et al. 2018), the activity of enhancers has not been investigated in sequential developmental stages, such as during worker metamorphosis. Because the worker caste evolved as the first sterile caste in eusocial bees such as *Apis mellifera* (Tian and Zhou 2014), its developmental process is an important target for understanding the molecular basis of caste evolution. During metamorphosis in *A. mellifera*, dramatic tissue remodeling results in increased cellular diversity and expansion in the number of genes expressed (Patir et al. 2023). Thus, worker metamorphosis provides an ideal model for investigating transcriptional regulation in *A. mellifera*. In holometabolous insects such as *A. mellifera*, the signaling pathways of juvenile hormone (JH) and ecdysone, mediated by *Methoprene-tolerant* (*Met*), *Krüppel-homolog 1* (*Kr-h1*), *Broad-complex* (*Br-c*), and *E93*, play central roles in the progression of metamorphosis (Belles and Santos 2014). A JH acid methyltransferase (JHAMT) is involved in JH biosynthesis in insects (Minakuchi et al. 2008; Goodman and Cusson 2012; Noriega 2014). *Met* functions as a JH receptor (Miura et al. 2005; Konopova and Jindra 2007; Charles et al. 2011; Jindra et al. 2015) and mediates the expression of *Kr-h1* (Minakuchi et al. 2009; Kayukawa et al. 2012). The active form of ecdysteroid, 20-hydroxyecdysone (20E), binds to the protein encoded by the ecdysone receptor gene (*EcR*) (Koelle et al. 1991; Yao et al. 1993) and induces the expression of *Br-c*, *E74*, and *E75*, which serve as ecdysteroid signaling mediators (Dubrovsky 2005). Expression analysis of these hormone signaling genes can be used to infer the endocrine status of individuals (Konopova et al. 2011). However, the regulatory relationships between these key metamorphosis genes and their enhancers remain largely unexplored in *A. mellifera*.

In this study, we used CAGE to identify active enhancers during worker metamorphosis in *A. mellifera*. Although CAGE has previously been applied to workers (nurses and foragers), its use has been limited to TSS identification, with no attempt having been made to detect enhancers (Khamis et al. 2015). Using these enhancers, we examined their regulatory relationships with the target genes. Furthermore, we investigated the sequence conservation of TFBSs within the identified enhancers and identified lineage-specific conservation patterns in bees.

## Results

### Landscape of CAGE Results

We performed CAGE across the developmental stages in worker metamorphosis, including larval, prepupal, and pupal stages and identified 17,349 unidirectional tag clusters representing candidate transcription start sites (TSSs), which were uniquely assigned to 8,535 genes. The unidirectional tag clusters were predominantly located in the promoter regions, consistent with the expected CAGE signal properties (Figure 1A). A total of 842 bidirectional tag clusters were identified. Among these, those located in the intronic or intergenic regions were considered enhancer candidates. This yielded 621 intronic and 221 intergenic enhancer candidates (Figure 1A).

**Fig. 1.**
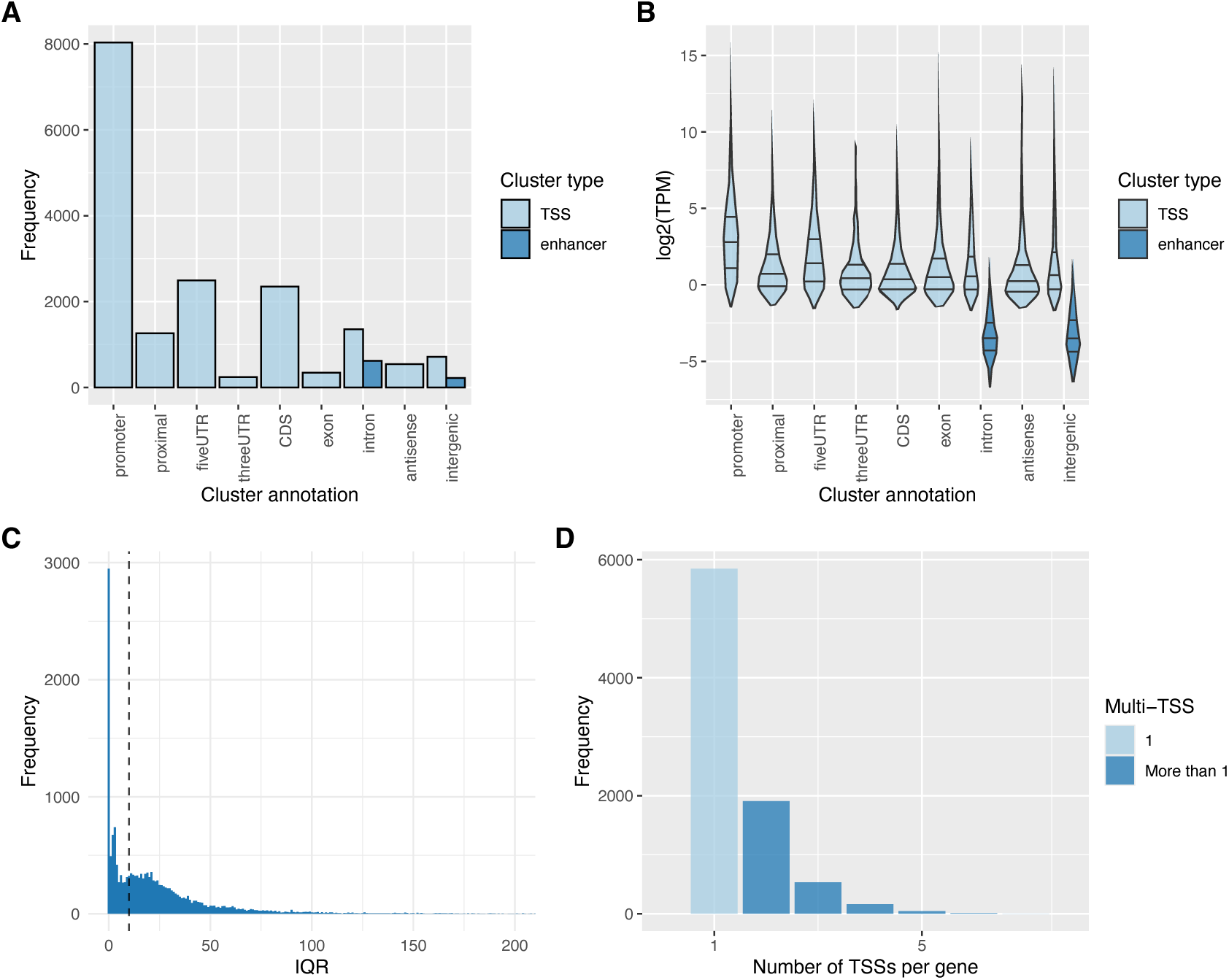
Overview of cap analysis of gene expression-sequencing (CAGE-seq) results. A) Annotation of tag clusters using RefSeq. Cluster types (transcription start site (TSS) or enhancer) are shown in different colors. B) Log2-scaled transcripts per million (TPM) values for each genomic region. Cluster types are indicated in different colors. C) Histogram of interquartile range (IQR) values for tag clusters. The dashed line marks an IQR value of 10 bp. D) Number of TSSs per gene. Light blue indicates genes with a single TSS, and dark blue indicates genes with two or more TSSs.

The expression levels of TSSs found in the promoter and 5′-untranslated regions (UTRs) were higher than of those found in the proximal regions (Figure 1B). Enhancer candidates consistently showed lower activity than the other regions (Figure 1B). TSSs, defined as collections of CAGE signals, were classified into two types: sharp and broad. The interquartile range (IQR) was used as an indicator of TSS width (Thodberg et al. 2019). The IQR values showed a bimodal distribution, with a dominant peak below 10 bp, indicating the predominance of sharp TSSs during the worker developmental stages (Figure 1C). Next, we examined the number of TSSs per gene and found that 68.7% (5,849/8,535) genes had a single TSS (Figure 1D). These results indicated that our CAGE data successfully captured transcriptional initiation and enhancer activity during worker metamorphosis, thereby providing a foundation for analyzing the regulatory relationships between TSSs and enhancers.

### Expression Patterns During Worker Metamorphosis

Using the k-means method incorporated into iDEP (Ge et al. 2018), we examined the expression patterns during worker metamorphosis and identified five distinct clusters (Figure 2A, Supplementary Table S1). These clusters were largely classified into two groups: those with high expression in larvae (days 9 and 11) and those with high expression in pupae (days 19 and 21). Principal component analysis revealed a clear separation by developmental stage, confirming that the identified expression patterns reflected progressive transcriptional changes during worker metamorphosis (Figure 2B).

**Fig. 2.**
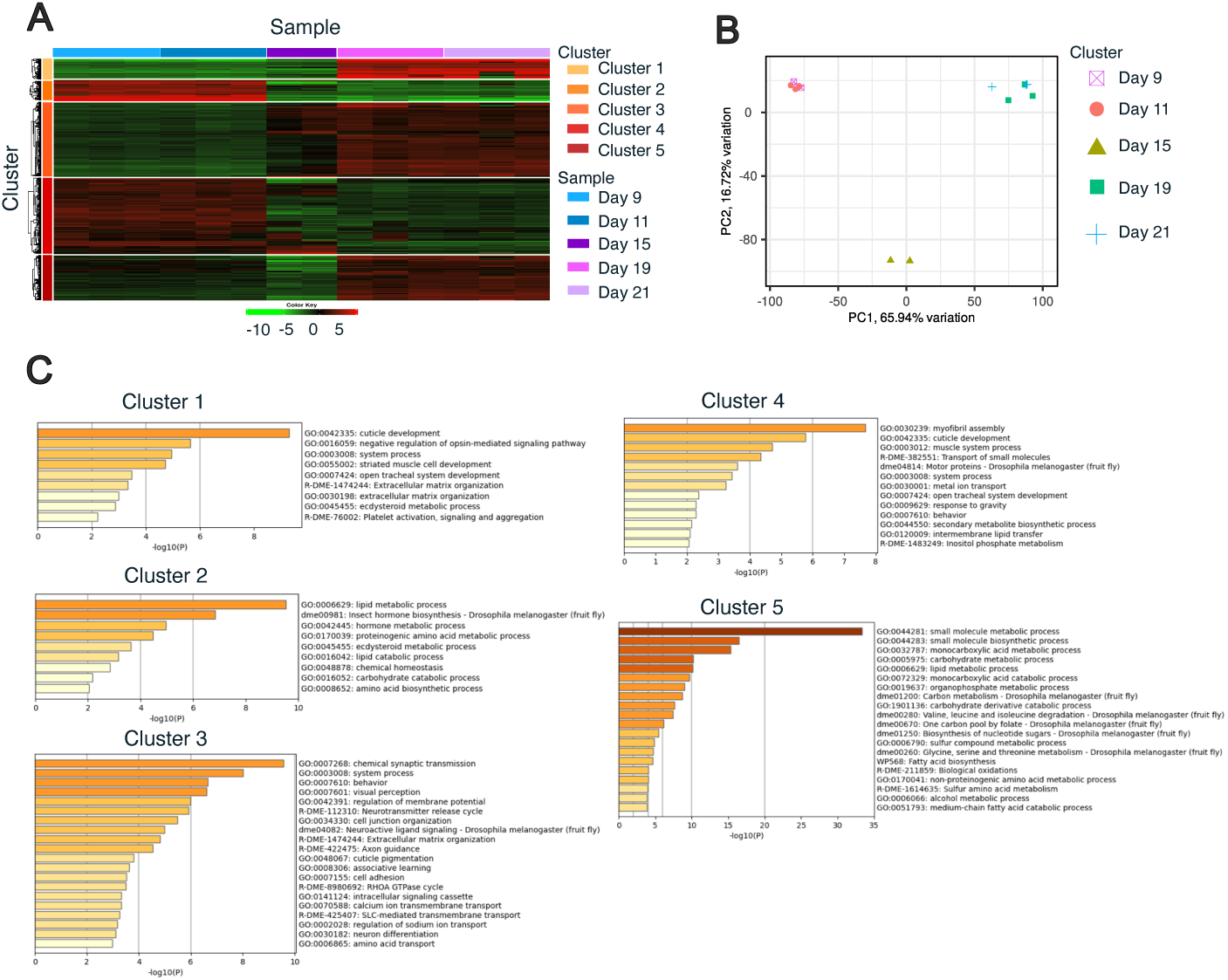
Gene expression dynamics during worker metamorphosis. A) Heatmap showing expression patterns across developmental stages. Red and green represent high and low expression levels, respectively. B) Principal component analysis (PCA) based on the top 2,000 genes. C) Metascape enrichment results. Bars represent −log10(*p*-value), and colors indicate the magnitude of enrichment. Gene ontology terms are shown on the right. To understand why *Kr-h1*, *E75*, and *EcR* were not assigned to any cluster, we compared our data with the published RNA-seq profiles (Yokoi 2024; Yokoi et al. 2025). *Kr-h1* showed the highest expression level on day 7 (larvae) (Figure 3), which is consistent with its role as an antimetamorphic factor. Although additional expression peaks were observed on days 11 and 13, our CAGE analysis began on day 9 and therefore did not capture the major peak on day 7. *Br-c* exhibited high expression levels between day 9 and 16, a period covered by our CAGE sampling. Therefore, *Br-c* was assigned to cluster 2. *E93* also showed high expression from day 13 to 19, which was broadly consistent with our CAGE data. In contrast, *E75* was not assigned to any expression cluster because RNA-seq data showed that its expression peaked on days 17 and 18, which were outside our CAGE sampling window. *EcR* was highly expressed on day 7 but did not show a distinct peak thereafter. Collectively, these results indicated that our CAGE data successfully captured key metamorphic regulators, such as *Br-c* and *E93*. The absence of *Kr-h1* and *E75* from the expression clusters can be explained by their expression peaks falling outside the sampling window, whereas *EcR* was not assigned, probably due to the lack of a distinct expression peak during the sampled stages.

To characterize the expression patterns of the five clusters, we performed an enrichment analysis using Metascape (Zhou et al. 2019) (Figure 2C and Supplementary Tables S2–S6). Each cluster was enriched for different gene ontology (GO) terms. Cluster 1 showed enrichment of the GO term “cuticle development”, which consisted of many genes encoding cuticle proteins (Supplementary Table S2). Cluster 2 was enriched for the “lipid metabolic process”. Careful examination of the gene list revealed LOC724386 (*Npc2a*) and LOC102655009 (*sit*) (Supplementary Table S3), which are involved in cholesterol trafficking and steroidogenesis (Huang et al. 2007; Danielsen et al. 2016). Cluster 3 was enriched for “chemical synaptic transmission” (Supplementary Table S4), indicating active neural remodeling during metamorphosis. Cluster 4 was enriched for the GO term “myofibril assembly” (Supplementary Table S5). Because the metamorphic process involves tissue remodeling, the enrichment of “myofibril assembly” in our dataset was consistent with expectations. Cluster 5 was enriched for the GO term “small molecule metabolic processes,” including LOC726445 (*Gpdh1*) and LOC411188 (*Ldh*) (Supplementary Table S6). These genes promote glycolytic flux that regulates larval growth (Li et al. 2019). Glucose metabolism plays a crucial role in pupal metamorphosis in *Drosophila melanogaster* (Nishimura 2020). Collectively, the enriched GO terms were consistent with known processes in insect metamorphosis, demonstrating that our CAGE data successfully captured the associated changes in gene expression during worker metamorphosis.

To verify whether the identified clusters captured changes in gene expression during metamorphosis, we selected five genes associated with metamorphic progression in insects (*Kr-h1*, ecdysone-induced protein 75 (*E75*), *Br-c*, *E93*, and ecdysone receptor (*EcR*)). We identified the expression clusters containing these genes (Figure 3A). *Br-c* and *E93* were included in Clusters 2 and 3, respectively, whereas *Kr-h1*, *E75*, and *EcR* could not be assigned to any expression clusters. *Br-c* was upregulated at the onset of prepupal stages (days 9 and 11), as expected for holometabolous insects (Truman 2019). *E93* expression levels increased at the onset of pupal stages (days 15, 19, and 21), consistent with the expression patterns observed for other insects (Belles and Santos 2014). The detection of these metamorphic regulators (*Br-c* and *E93*) indicated that transcriptional changes during worker metamorphosis were successfully captured in our CAGE data.

**Fig. 3.**
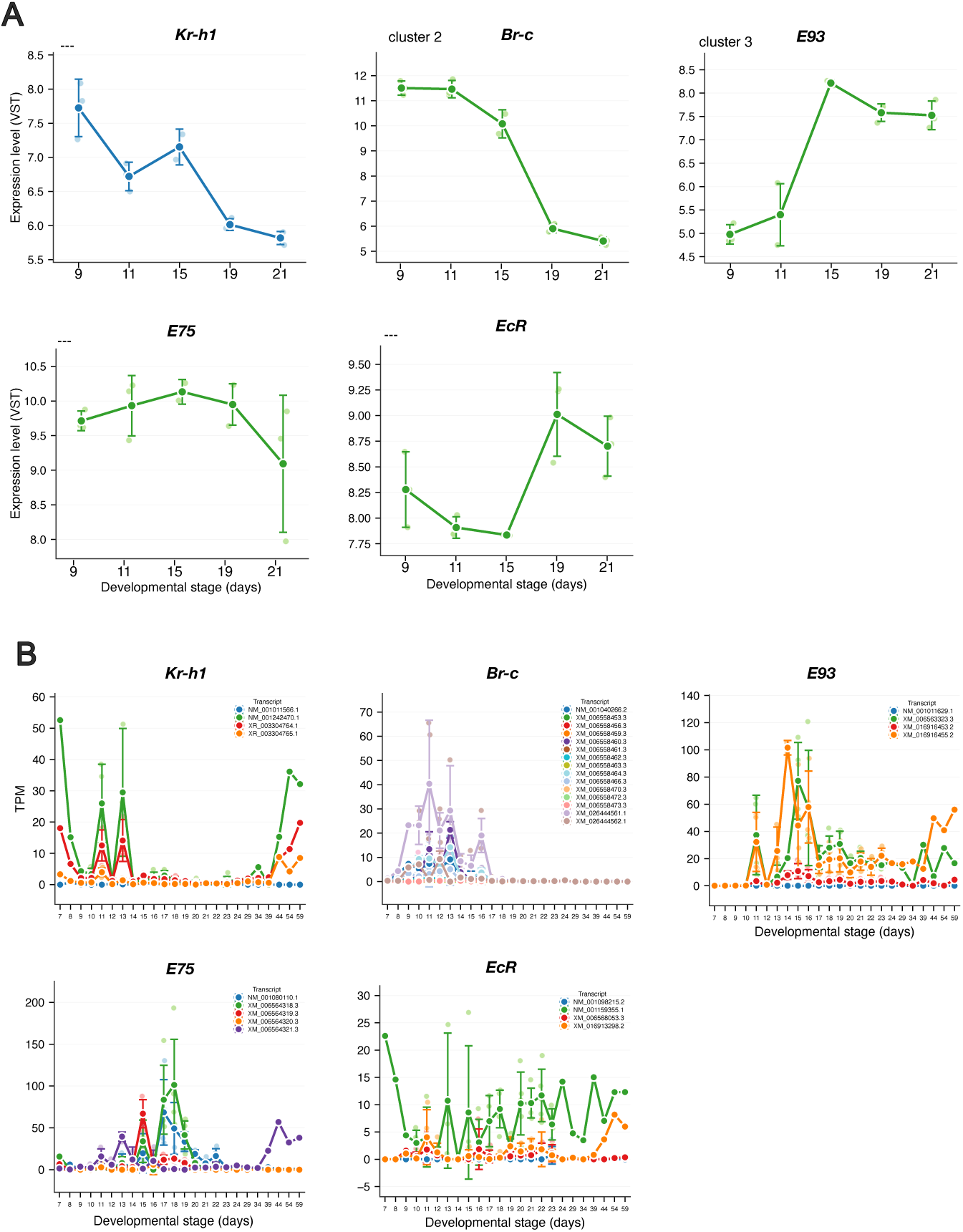
Expression patterns of metamorphosis-related marker genes. A) Expression patterns of metamorphic marker genes in our cap analysis of gene expression (CAGE) dataset. Blue and green plots represent genes responsive to juvenile hormone and ecdysone, respectively. Variance stabilizing transformation (VST)-normalized counts are shown. The expression clusters identified in this study are indicated in the top left of each panel; “---” indicates genes not assigned to any cluster. B) Expression patterns throughout worker development based on published RNA sequencing data (Yokoi et al. 2025). Transcripts per million (TPM) values for all transcripts of each gene are shown. Larvae corresponded to days 6–9 post-oviposition, prepupal and pupal stages to days 10–18, and adults to day 19 and later.

### Identification of the Transcriptional Regulation During Worker Metamorphosis

We examined the relationships among transcription factors (TFs), enhancers (E), and TSSs to identify transcriptional regulation during worker metamorphosis. Using the CAGEfightR “findLinks” program, we identified a total of 1,505 E-TSS pairs with correlated expression level (*r* > 0, *p* < 0.05, Supplementary Table S7). Because genes within the same cluster exhibited similar expression patterns (Figure 2A), we hypothesized that they were regulated by common TFs. To test this hypothesis, we performed motif enrichment analysis to identify common TFBSs within each cluster using simple enrichment analysis (SEA) in the MEME suite (Supplementary Table S8–S12). Additionally, we examined the correlation of expression levels between the identified TFs and TSSs using our CAGE-seq data (*p* < 0.05, Spearman’s rank correlation coefficient >0; Supplementary Table S13). By integrating these results, we identified the TF–E–TSS relationships within each cluster, resulting in 15 sets of these relationships (Table 1). Among these 15 sets, *tramtrack* (*ttk*) was the transcription factor associated with the largest number of target genes, including the metamorphic regulator *Br-c* (Table 1). The predicted enhancers for *Br-c* were located within introns (Figure 4A and 4B, Br-c_enhancer_1: NC_037650.1:3332653–3333205; Figure 4A and 4C, Br-c_enhancer_2: NC_037650.1:3355266–3355678) (Table 1). Br-c_enhancer_1 activity correlated with seven TSS activities observed in the range NC_037650.1:3337344–3342410 (Figure 4A arrows and Supplementary Table S7), whereas Br-c_enhancer_2 activity correlated with a single TSS activity in the range NC_037650.1:3358628–3358852 (Figure 4A arrowhead and Supplementary Table S7).

**Fig. 4.**
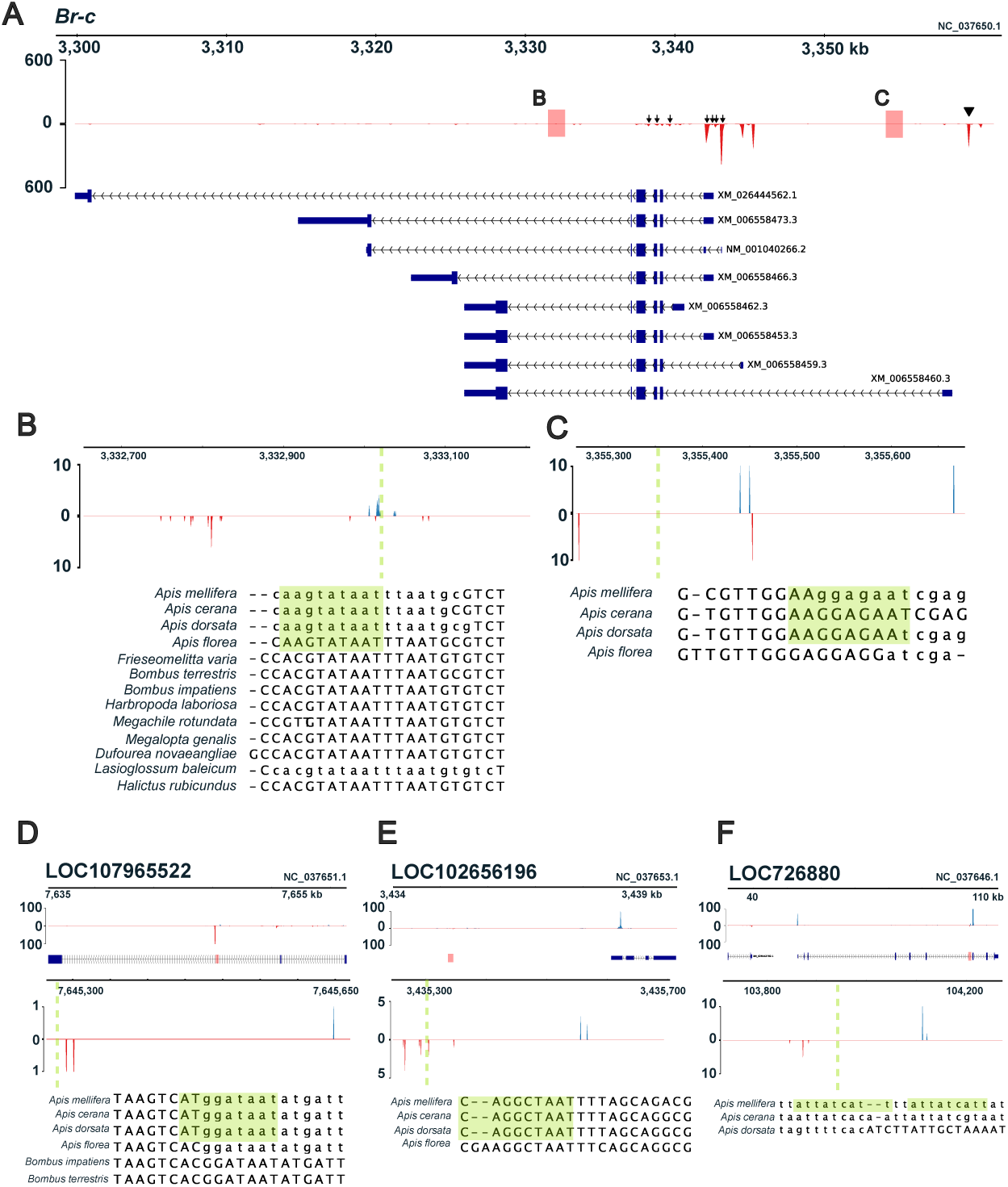
Enhancer regions containing the *tramtrack* (*ttk*)-binding sites. In all the figure panels, red and blue peaks indicate signals on the negative and positive strands, respectively. Green boxes and dashed lines indicate predicted *ttk*-binding sites (from *Drosophila melanogaster* motifs). Gene models are shown in blue. Corresponding enhancer regions in other bee species are displayed at the bottom of each panel. Lowercase nucleotide sequences represent soft-masked repeat regions in the genome. The y-axis represents the cap analysis of gene expression transcriptional start site (CTSS) count data. (A) Tag clusters and gene structure for *Br-c*. Arrowheads and arrow indicate each TSS. Red boxes represent enhancer regions identified in this study; regions B and C are magnified in panels B and C. Arrows and arrowheads indicate transcription start sites. (B) Intronic enhancer region in *Br-c* (magnified view of region B in panel A). (C) Additional intronic enhancer region in *Br-c* (magnified view of region C in panel A). (D) Intronic enhancer region of LOC107965522. (E) Intergenic enhancer region of LOC102656196. (F) Intronic enhancer region of LOC726880.

**Table 1.**
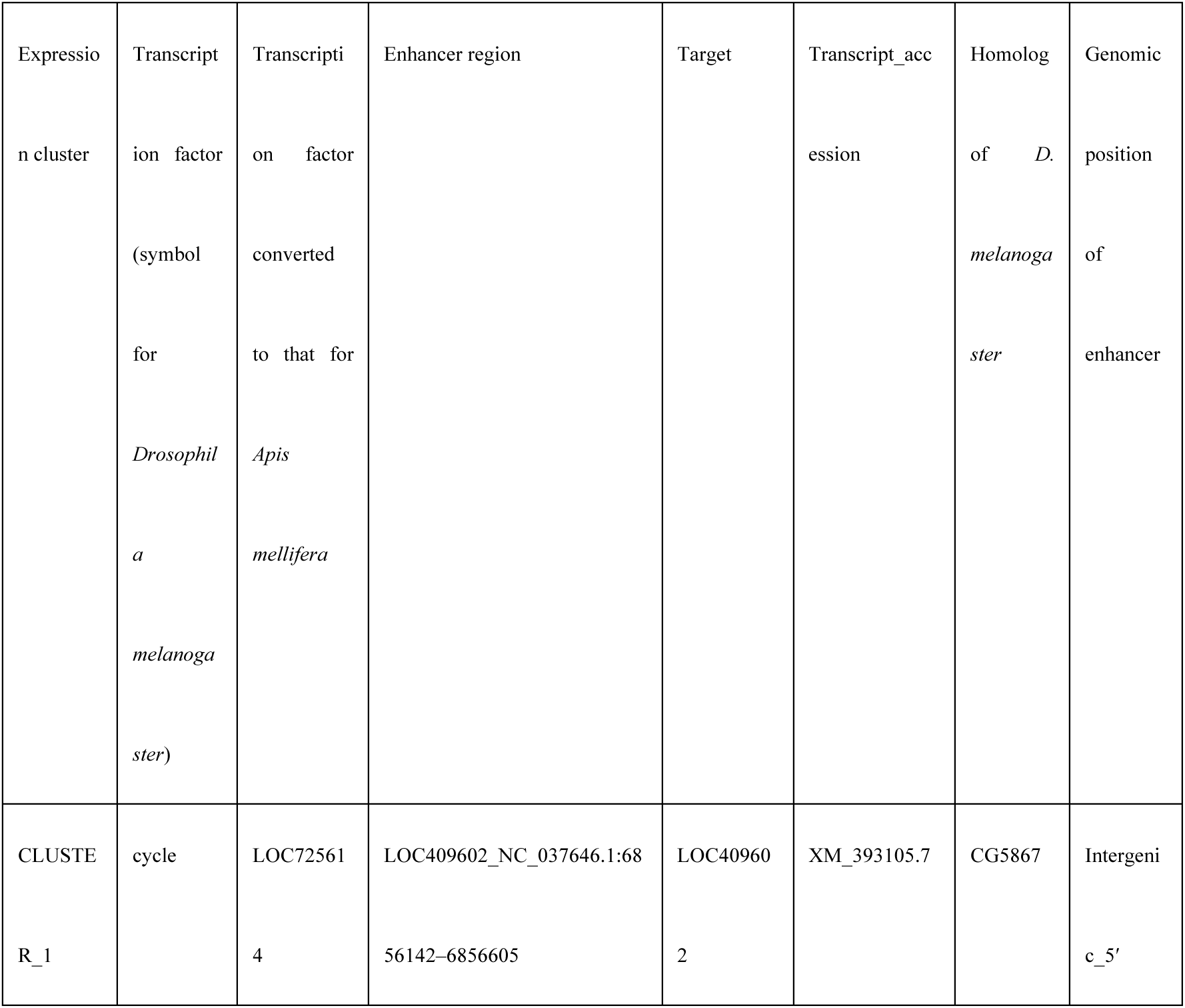
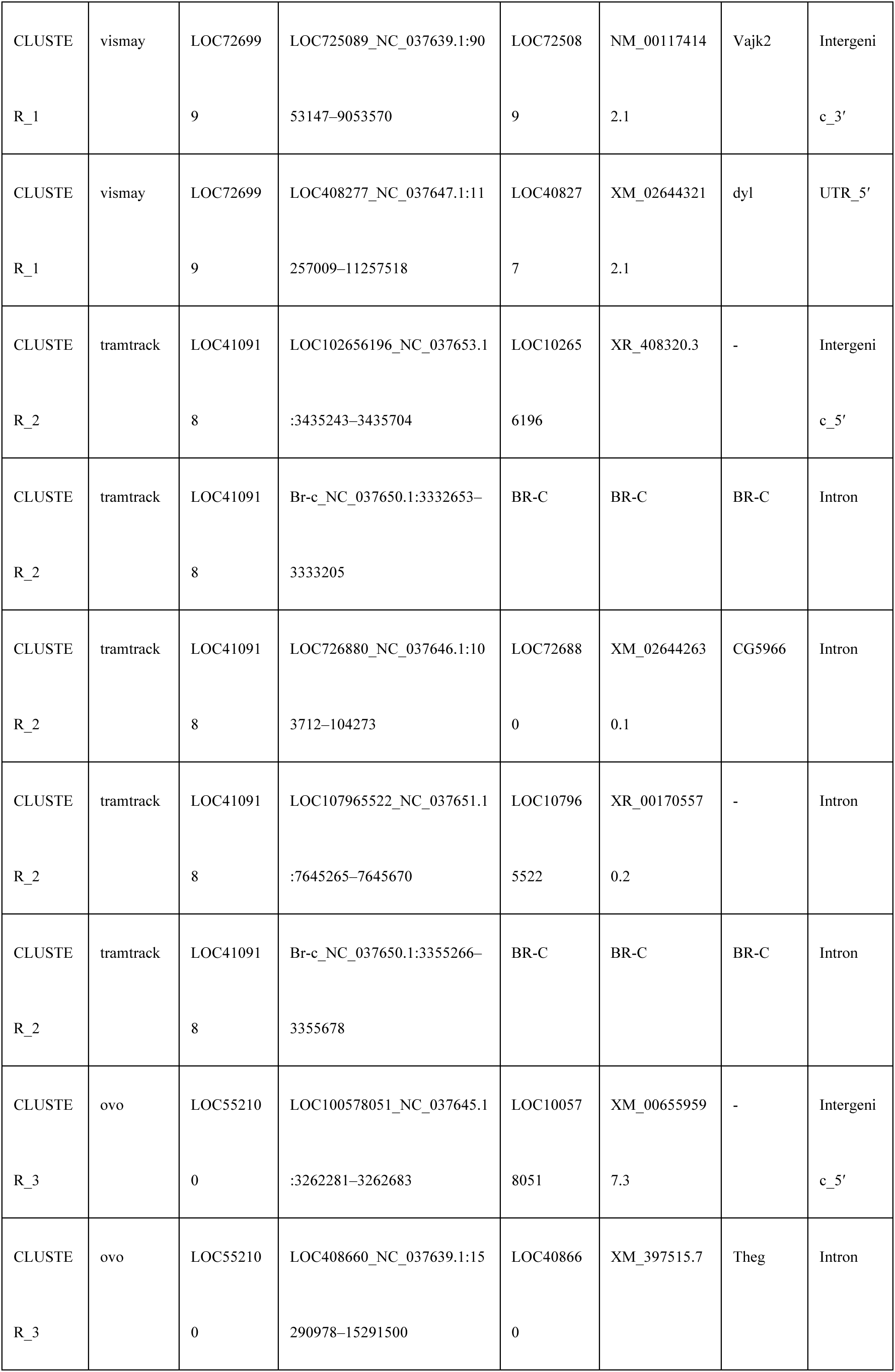
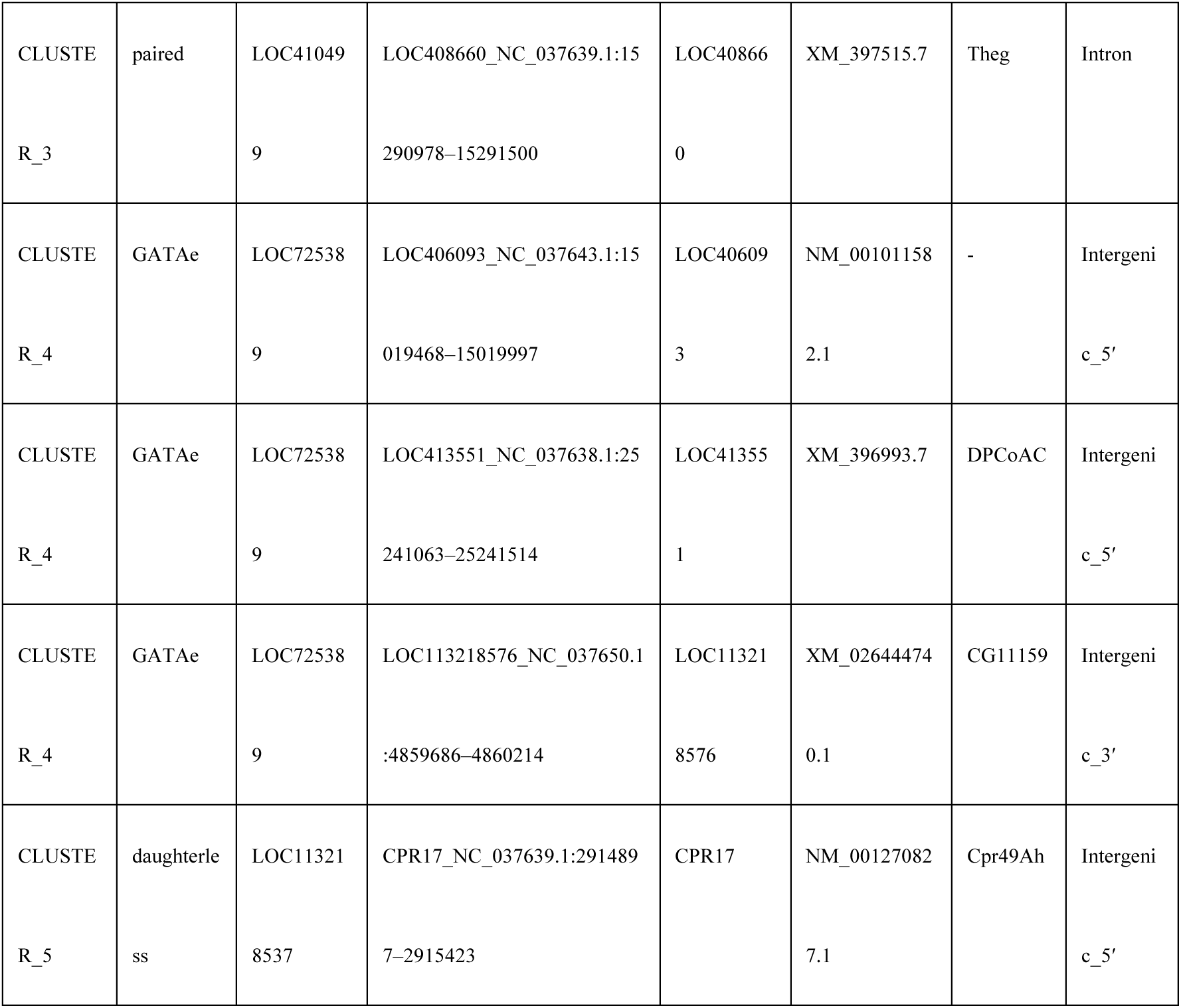
Summary of transcription factors, enhancer regions, and corresponding target genes identified in this study.

Next, we examined the sequences of *ttk*-binding sites in Br-c_enhancer_1 and _2. SEA identified the *ttk*-binding site to have a consensus sequence: MAGTATAAT (Supplementary Table S9). In Br-c_enhancer_1, we identified the *ttk*-binding site as AAGTATAAT (Supplementary Table S14). The *ttk*-binding site in Br-c_enhancer_1 was located near one of the bidirectional tag clusters (Figure 4B, green dashed line). The identified *ttk*-binding sites (AAGTATAAT) were conserved only within the *Apis* genus (Figure 4B, green box). Almost all other bee species had the same sequence, ACGTATAAT, in their corresponding region, except for *Megachile rotundata*. In Br-c_enhancer_2, the *ttk*-binding site (AAGGAGAAT, Supplementary Table S14) was located downstream of the *Br-c* TSS (Figure 4C, green dashed line). Because of low sequence conservation, the regions corresponding to the *ttk*-binding site in Br-c_enhancer_2 could not be identified in non-*Apis* species. Within the *Apis* genus, the binding site was conserved across species, except for *Apis florea* (Figure 4C, green box).

We further investigated the sequence conservation of the *ttk*-binding sites in other target genes (Table 1), including LOC107965522 (ncRNA), LOC102656196 (ncRNA), and LOC726880 (pancreatic lipase-related protein 2-like) (Figure 4D–F). Unlike LOC107965522 and LOC726880, the enhancer of LOC102656196 was located 3 kb upstream of the TSS (Figure 4E). Sequence conservation of all *ttk*-binding sites was limited to the *Apis* genus (Figure 4D and 4E) and *A. mellifera* (Figure 4F). As observed for Br-c_enhancer_1, the *ttk*-binding sites of LOC107965522 and LOC102656196 were located near the enhancer tag clusters (Figure 4D and 4E, green dashed lines), whereas those of LOC726880 did not overlap with the enhancer tag clusters (Figure 4F, green dashed line).

## Discussion

### Genome-wide Detection of TSSs and Enhancers During Worker Metamorphosis

We identified genome-wide TSS activity during worker metamorphosis in *A. mellifera*. Most CAGE tag clusters were detected as TSSs in the promoter regions of reference gene annotations. Additionally, 842 enhancers were identified, of which intronic enhancers were more prevalent than intergenic enhancers. While previous searches for TFBSs based on sequence alone focused on the 5′ upstream region of genes (Kapheim et al. 2015; Shell et al. 2021), our results revealed numerous intronic enhancers. This finding is consistent with a previous study demonstrating that active intronic enhancers are abundant in worker larvae (Wojciechowski et al. 2018). In humans, intronic enhancers predominantly regulate tissue-specific expression of genes (Borsari et al. 2021). As worker metamorphosis involves tissue remodeling toward adult tissue development, the intronic enhancers identified in this study may contribute to tissue-specific expression of genes. Evidence of tissue-specific expression was also reflected in the shape of the TSS clusters, as assessed via the IQR of tag positions. Sharp TSSs (i.e., IQR < 10) occurred more frequently than broad TSSs. Sharp TSSs are predominantly associated with tissue-specific activity in mammals (Carninci et al. 2006). Collectively, our CAGE data highlighted the importance of tissue-specific gene expression during worker metamorphosis.

### Identification of Active TSS During Worker Metamorphic Process

Five clusters were identified based on the expression patterns. Enrichment analysis revealed that each cluster was comprised of genes with distinct biological functions. These GO terms were associated with insect metamorphosis, which indicated that the changes in expression detected using CAGE reflect the developmental transition during metamorphosis in *A. mellifera* workers. This finding is supported by the dynamic changes in the expression of *Br-c* and *E93*, which are regulators of metamorphosis. However, some known marker genes (*Kr-h1*, *E75*, and *EcR*) did not show significant changes in their expression. For *Kr-h1* and *E75*, this was probably because of the fact that our sampling covered only a subset of the developmental stages, and their expression peaks occurred outside the stages sampled in this study. *EcR* is generally involved in metamorphic signaling in insects; however, we did not detect distinct changes in its expression in our dataset. This is consistent with previous findings regarding no distinct expression pattern of *EcR* in wing disks during *A. mellifera* metamorphosis (Soares et al. 2021). These results suggest that the expression dynamics of *EcR* may differ between tissues or that its regulatory role in worker metamorphosis is more complex than that in other insects.

### Regulatory Interaction Between *ttk* and *Br-c*

To identify the transcription factors that coordinately regulate gene expression within each cluster, we searched for TFBSs enriched in the enhancers of genes belonging to each expression cluster. Among the predicted target genes, *Br-c* was identified in the resulting list (Table 1). In this study, *ttk* was predicted to bind to the largest number of target genes, including *Br-c*. *ttk* is a member of the C2H2 zinc finger transcription factor family. Previous transcriptome analyses have shown that *Br-c* is more highly expressed in the wing disk during worker metamorphosis than during queen metamorphosis in *A. mellifera* (Soares et al. 2021). This finding supports the potential role of *ttk* in the upregulation *of Br-c* expression during worker metamorphosis.

The *ttk*-binding sites were located predominantly on one side of the enhancers rather than at their center. In humans, TFBSs are not always located at the center of enhancers, but are often found near eRNA TSSs (Grossman et al. 2018). In addition, the balance of bidirectional eRNA activity is associated with target gene expression in humans (Kristjánsdóttir et al. 2020). Similarly, *ttk*-binding sites overlapping eRNA TSSs may modulate the balance of eRNA activity in honeybees.

### Conservation of *ttk*-binding Sites and Eusocial Evolution

The number of TFBSs for a particular TFs tends to increase with the correlation of social complexity (Kapheim et al. 2015; Shell et al. 2021). *ttk* was classified as a TF related to social complexity, although this classification was based on findings in stingless bees and not in honeybees (Kapheim et al. 2015). We searched for sequence conservation of the *ttk*-binding site to determine whether common mechanisms regulating *Br-c* expression are present in bees. However, the conservation of the *ttk*-binding site did not correlate with eusocial complexity, and sequence conservation was observed only within the genus *Apis*. Kapheim et al. (2015) noted this inconsistency, showing that eusociality has evolved through different mechanisms across lineages. Our results are consistent with these findings. Nevertheless, *ttk* is a TTK-type BTB protein that forms hexameric complexes composed of different transcription factors, which is suggestive of diverse binding site compositions (Bonchuk et al. 2024). Therefore, further CAGE and chromatin immunoprecipitation-sequencing analyses in species other than *A. mellifera* are required to examine the precise and active *ttk*-binding sites.

### Conclusions and Study Limitations

In this study, we performed CAGE-seq during worker metamorphosis in *A. mellifera* to identify the regulatory elements and their interactions that control gene expression. We identified the promoters of 8,535 genes and 842 candidate enhancers. Furthermore, we identified *ttk* as a transcription factor that potentially regulates *Br-c* expression, with *ttk*-binding site conservation observed only for the genus *Apis*. However, the lack of sequence conservation outside the *Apis* genus may not necessarily indicate the absence of *ttk*-mediated regulation, because transcription factors can recognize diverse binding sequences. Therefore, integrating assay for transposase-accessible chromatin using sequencing, chromatin immunoprecipitation-sequencing, and chromosome conformation capture across bee species is necessary to fully elucidate enhancer function and validate the identified regulatory interactions. Additionally, the identified enhancers could be candidate targets for genome editing to validate their biological functions.

## Materials and Methods

### Honeybee Rearing and Sample Preparation

*A. mellifera* from an Italian hybrid strain reared at the apiary of the National Agricultural Research Organization (NARO) were used. *A. mellifera* was reared following standard beekeeping practices (Yokoi et al. 2025). Briefly, the queens were confined to egg-laying boxes for 6 h to obtain age-controlled samples. *A. mellifera* workers (days 6–58) were collected, with two to three biological replicates per time point. Samples were stored at −80 °C until use. For CAGE, RNA was extracted from larvae (days 9 and 11), prepupae (day 15), and pupae (days 19 and 21). This study involved only invertebrate animals (*Apis mellifera*), and therefore ethical approval was not required according to institutional and national guidelines.

### RNA Extraction

Total RNA was extracted from whole-body samples using TRIzol Reagent (Thermo Fisher Scientific) and purified with an RNeasy Mini Kit (Qiagen), according to the manufacturer’s protocols. The RNA concentration was measured using a NanoDrop spectrophotometer (Thermo Fisher Scientific). One to five biological replicates per sample category were used for RNA-Seq based on RNA yield and quality.

### CAGE Sequencing and Data Analysis

CAGE libraries were prepared by DNAFORM (Japan). Raw CAGE-seq reads were filtered using fastp v 1.0.1 (Chen et al 2025), and mapped to the *A. mellifera* genome (Accession ID: GCF_003254395.2) using STAR with default settings. BAM files were converted to bigwig files within CAGEfightR (Thodberg et al 2019), and these bigwig files were loaded into CAGEfightR to identify the CAGE transcriptional start site (CTSS). CTSSs detected in only a single sample were removed. The CTSSs were classified into unidirectional (TSS candidates) and bidirectional (enhancer candidates) clusters. Unidirectional clusters were filtered by >1 TPM for at least two samples. Bidirectional clusters obtained from at least three samples were filtered by balance threshold score >0.95. To convert HAv3.1 of *A. mellifera* (GCF_003254395.2 and GCF_003254395.2_Amel_HAv3.1.gff) into the BSgenome data package, BSgenomeForge v 1.74.0 (https://bioconductor.org/packages/release/bioc/html/BSgenomeForge.html, accessed on 8 April, 2025) was used. The converted BSgenome was used for gene-level annotations.

### Clustering of Gene Expression

iDEP v 2.01 (Ge et al. 2018), launched using Docker according to GitHub (https://github.com/gexijin/idepGolem?tab=readme-ov-file), was used for the expression analysis of unidirectional clusters. The count data of unidirectional clusters were loaded into iDEP and normalized using the variance stabilizing transformation (VST) method. Normalized count data were used for k-means clustering. The k-means clustering was performed for the top 2000 genes, and the number of clusters was set to five in the iDEP setting. Principal component analysis was performed using PCAtools in iDEP. To characterize the gene set, including expression clusters identified by k-means clustering, enrichment analysis was performed using Metascape (Zhou et al. 2019) with annotations identified previously (Yokoi et al. 2022). VST-normalized count data were used to visualize the expression patterns of individual genes.

### Identification of the TF–Enhancer–Gene Relationship

Correlation analysis between enhancer activity and gene expression was performed using the Kendall’s rank correlation coefficient with the findlinks function in CAGEfightR within a distance of 10k bp. Pairs of enhancer–gene interactions were filtered based on a positive correlation and *p*-value <0.05. TF-binding sites were searched using SEA in the MEME Suite 5.5.9, with sequences of biding sites of *D. melanogaster* from JASPAR (Ovek Baydar et al. 2026) (https://jaspar.elixir.no/downloads/ accessed on 22 Sep, 2025). Spearman’s rank correlation coefficient was calculated to evaluate the relationship between transcription factor and target gene expression patterns across developmental stages. Statistical analysis was performed using Python (version 3.12.7) with the SciPy library (version 1.14.1). Correlations with *p*-value <0.05 were considered statistically significant.

### Evaluation for Conservations of Sequences of TF-biding Sites

Conservation of the identified TF-binding sites was evaluated via DNA sequence alignments using a Threaded Blockset Aligner (TBA) (Blanchette et al. 2004). For this analysis, megablast using enhancer sequences was conducted against the RefSeq Genome Database (refseq_genomes) in Apidae (taxid: 7458), Megachilidae (taxid: 124286), and Halictidae (taxid: 77572) (https://blast.ncbi.nlm.nih.gov/Blast.cgi?PROGRAM=blastn&PAGE_TYPE=BlastSearch&LINK_LOC=blasthome, accessed on 19 January, 2026) to preliminarily identify the corresponding chromosome or contigs in other bees. In the TBA analysis, the phylogenetic relationships in bees were based on previous studies (Kapheim et al. 2015; Da Silva 2021). Expression peaks obtained from CAGE-seq were visualized using pyGenomeTracks version 3.9 (Lopez-Delisle et al. 2021).

## Acknowledgments

This work was supported by the Center of Innovation for Bio-Digital Transformation (BioDX), an open innovation platform for industry-academia co-creation (COI-NEXT) of JST (JPMJPF2010) to K.Y. and H.B. We thank all laboratory members at Hiroshima University for their valuable comments. Computations were performed on the computers at the Hiroshima University Genome Editing Innovation Center. Also, we thank Drs. Masatsugu Hatakeyama and Kiyoshi Kimura for preparing *A. mellifera* samples.

## Data Availability

The Raw CAGE data underlying this article are available in the DDBJ Sequence Read Archive at [https://www.ddbj.nig.ac.jp/], and can be accessed with the accession numbers DRR893670–DRR893683. All supplementary results are available at figshare (https://doi.org/10.6084/m9.figshare.31237126, accessed on 4 Mar 2026) as follows. Supplementary Table S1: Gene expression clusters identified via k-means analysis; Supplementary Table S2: Gene ontology (GO) enrichment using Metascape for Cluster 1; Supplementary Table S3: Gene ontology (GO) enrichment using Metascape for Cluster 2; Supplementary Table S4: Gene ontology (GO) enrichment using Metascape for Cluster 3; Supplementary Table S5: Gene ontology (GO) enrichment using Metascape for Cluster 4; Supplementary Table S6: Gene ontology (GO) enrichment using Metascape for Cluster 5; Supplementary Table S7: Correlation analysis between enhancer activity and transcription start site (TSS) expression; Supplementary Table S8: Simple enrichment analysis (SEA) motif enrichment for Cluster 1; Supplementary Table S9: Simple enrichment analysis (SEA) motif enrichment for Cluster 2; Supplementary Table S10: Simple enrichment analysis (SEA) motif enrichment for Cluster 3; Supplementary Table S11: Simple enrichment analysis (SEA) motif enrichment for Cluster 4; Supplementary Table S12: Simple enrichment analysis (SEA) motif enrichment for Cluster 5; Supplementary Table S13: Correlation analysis between transcription factor expression and transcription start site (TSS) activity; Supplementary Table S14: ttk-binding site sequences identified in the enhancer regions.

## Author Contributions

Conceptualization, K.Y. and H.B.; methodology, K.T. and H.B.; software, K.T.; validation, K.T., K.Y., and H.B.; formal analysis, K.T.; investigation, K.Y.; resources, K.Y. and H.B.; data curation, K.T.; writing—original draft preparation, K.T. and K.Y.; writing—review and editing, K.T., K.Y., and H.B.; visualization, K.T.; supervision, H.B.; project administration, H.B.; funding acquisition, K.Y. and H.B. All authors have read and agreed to the published version of the manuscript.

